# Text mining of CHO bioprocess bibliome: Topic modeling and document classification

**DOI:** 10.1101/2022.08.22.504864

**Authors:** Qinghua Wang, Jonathan Olshin, K. Vijay-Shanker, Cathy Wu

**Author notes:** Corresponding author, (QW). These authors contributed equally to this work (Qinghua Wang, Jonathan Olshin).

## Abstract

Chinese hamster ovary (CHO) cells are widely used for mass production of therapeutic proteins in the pharmaceutical industry. With the growing need in optimizing the performance of producer CHO cell lines, research on CHO cell line development and bioprocess continues to increase in recent decades. Bibliographic mapping and classification of relevant research studies will be essential for identifying research gaps and trends in literature. To qualitatively and quantitatively understand the CHO literature, we have conducted topic modeling using a CHO bioprocess bibliome manually compiled in 2016, and compared the topics uncovered by the Latent Dirichlet Allocation (LDA) models with the human labels of the CHO bibliome. The results show a significant overlap between the manually selected categories and computationally generated topics, and reveal the machine-generated topic-specific characteristics. To identify relevant CHO bioprocessing papers from new scientific literature, we have developed a supervised learning model, Logistic Regression, to identify specific article topics and evaluated the results using three CHO bibliome datasets, Bioprocessing set, Glycosylation set, and Phenotype set. The use of top terms as features supports the explainability of document classification results to yield insights on new CHO bioprocessing papers.

## 1. Introduction

### CHO biblioem

Chinese hamster ovary (CHO) cells are widely used for biological and medical research (1, 2). They are the predominant host for mass production of many therapeutic proteins such as recombinant monoclonal antibodies in the pharmaceutical industry (3). With the increasing market demand and growing need in optimizing the performance of producer CHO cell lines, research on CHO cell line development and bioprocess engineering continuously increases in recent decades (4, 5). In 2016, Golabgir et al. (6) reported a manual bibliographic compilation of the published CHO cell studies from January 1995 to June 2015, which were retrieved with keywords “CHO cells” and/or “Chinese hamster ovary” in the title or abstract from Thomson Reuters Web of Science™ The initial article set (10,279 abstracts) was manually filtered to identify a bioprocess (BP) set (1157 abstracts) that focus on CHO cell bioprocesses and biotechnologies, including host cell line engineering, strain selection/screening, and cell culture media design, etc. The non-BP set covers the remaining abstracts describing studies irrelevant to CHO bioprocess. For each BP abstract in the CHO bibliome, one or more category labels from a total of 16 research categories were manually assigned based on the types of phenotypic and bioprocess data contained therein (6).

The CHO bibliome continues to grow since its last compilation in 2015, with over 500 PubMed citations annually. To automate text analysis of the CHO bibliome and gain insight to key topics and trends in CHO bioprocessing and biotechnologies, we have applied topic modeling to explore and classify CHO literature and compared results with those manually assigned category labels in the CHO bibliome. When coupled with our classifiers trained with supervised machine learning methods, the resulting models can automatically classify the newly published CHO cell studies after 2015 into bioprocess categories and help researchers select CHO cell research articles of their interest.

### Topic modeling and document classification

Natural language processing (NLP) allows machines to interpret human language with either unsupervised or supervised approaches (7, 8). For text analysis to uncover the main topics in an unlabeled set of documents, probabilistic topic models are considered an effective framework for unsupervised topic discovery (9, 10). Latent Dirichlet Allocation (LDA) is a widely used topic modeling method (11) with many applications (12). It is a generative probabilistic model of a corpus. The basic principle is that documents are represented as random mixtures over latent (hidden) topics, where each topic is characterized by a distribution over words in the corpus. In this study, LDA is adopted for an automatic exploration of latent topics in the CHO bioprocess bibliome, which are then compared and contrasted with those previously manually assigned research categories. This allows to gain insight into practical performance of LDA topic models in comparison with human manual category labels, and potential benefits from applying topic modeling to identify significant topics.

To identify new CHO bioprocessing papers from PubMed (especially for publications after 2015), a classifier is needed to separate BP from non-BP studies and identify their bioprocessing topics by learning how the existing CHO bibliome classifies them. For this task, a supervised approach, Logistic Regression, is utilized to classify the bibliome using three datasets, one for the overall “Bioprocess” category (BP set), and two on the specific bioprocessing categories of “Phenotype and Production Characteristics” (Phenotype set) and “Glycosylation” (Glycosylation set), respectively. Logistic regression allows for different term representations to be used in classification efficiently, ranging from a term’s binary presence/absence method, term frequency (tf), and term frequency-inverse document frequency (tf-idf) (13). Our objective is to determine if each category of interest includes unique terms that could be used for document classification. If the model is able to predict the category of a document in a dataset with high accuracy, it suggests that the documents in that category share an adequate amount of unique terms for classification, which may yield insights on new CHO bioprocessing papers.

## 2. Methods

The CHO bibliome processing and analysis workflow consists of document processing, unsupervised topic modeling, and supervised document classification, as summarized in Fig 1.

**Figure 1.**
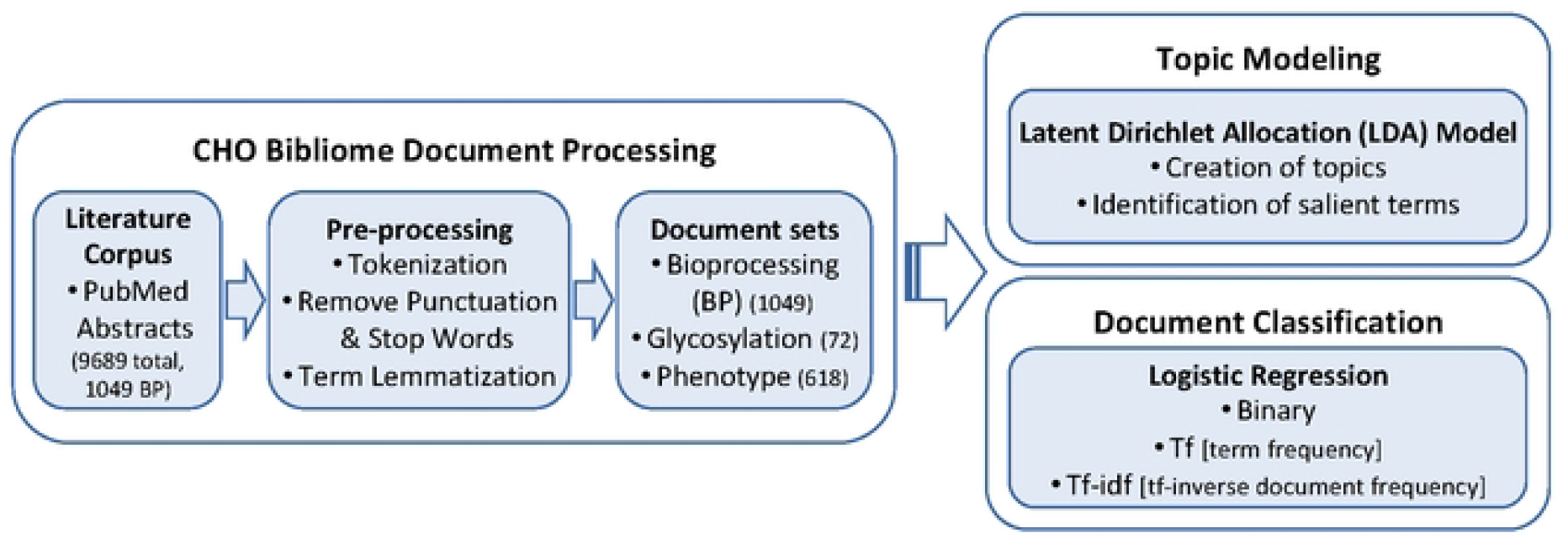
Overview of the CHO bibliome processing and analysis workflow.

### 2.1 Document processing

#### Literature corpus

For both topic modeling and document classification tasks in this study, we used the abstracts that were compiled in the CHO bibliome paper (6). To retrieve the abstract texts for the citations in the bibliome, PubMed was used for obtaining PMIDs based on matching of title, doi, and/or journal information. PubTator API was used to retrieve the documents containing both title and abstract text with PMID as query (14). The resulting dataset consisted of 9689 documents, including 1049 documents in the Bioprocessing (BP) set and 8640 documents in the non-BP set.

#### Pre-processing

The text processing included typical NLP steps: removal of special characters and numbers, removal of stop words (NLTK package (15)), tokenization, and term lemmatization (with Part of Speech (POS) Tagging allowed for ‘NOUN’, ‘VERB’, ‘ADJ’, ‘ADV’,’PROPN’, ‘NUM’; with spaCy library (13, 16)). The processing was conducted within a Jupyter notebook (17).

Further text processing for document classification was conducted within a Jupyter notebook using Python. The dataset of 9689 documents (BP + non-BP) were mapped to their designated topic classification as marked in (6). The articles were labeled with a 0 or 1 signifying each document’s allocation to the negative and positive set, respectively, for each bioprocess category of interest. These articles with known bioprocess classification allow a training set and test set to be created. 5-fold cross-validation was used for checking the accuracy of the model, where 80% of the dataset was used as the training set and 20% as the test set each time.

#### Document sets

Three document sets were compiled to study the efficiency and accuracy of the classifiers for predicting previously unseen documents. The BP set consisting of 1049 BP and 8640 non-BP documents was used to discern bioprocessing-related papers from all CHO cell publications. Two human-labeled categories of the BP documents, Glycosylation and Phenotype, were used to study how well the system can automatically identify articles containing these two bioprocess categories of interest. The Glycosylation set consisted of 70 Glycosylation and 979 non-Glycosylation BP documents, while the Phenotype set consisted of 547 Phenotype and 502 non-Phenotype BP documents.

### 2.2 Topic modeling using LDA

LDA is among the most widely applied probabilistic topic modeling approach (12). Python’s GENSIM package(18) was used for LDA applications in this study. Bigrams and trigrams were created with GENSIM phrase detection and added to the dictionary. Words that appear in less than 5 documents were filtered out, resulting in 2534 words in the final dictionary. Lastly, BP documents were included in training the LDA model with Python’s GENSIM package. Method of grid search is employed to select the best set of hyperparameters (i.e., the number of topics, alpha, and eta) for the final LDA model. The resulting document-to-topic probabilities based on the chosen model were analyzed and compared with the previously reported manual category assignments. Python library pyLDAvis (19) is used for interactive topic model visualization.

### 2.3 Document classification

#### Logistic regression

It was implemented for classification of three document sets, BP, Glycosylation, and Phenotype. Multiple trials were conducted. The first trial was run using a binary term representation feature with the entirety of the vocabulary. The binary feature specifying whether a term was present or not in the article was considered. The results thus represented as a baseline for the performance of the logistic regression on classification tasks.

The second trial involved the use of tf-idf (13), which is often used to capture the importance of the terms in the document. Only terms with minimum document frequency (df_min) of 0.05 and a maximum document frequency (df_max) of 0.95 were used. This allowed words to be removed that only occurred a very limited amount of times in the dataset. The use of the entire feature list allows the feature list length to be altered in subsequent trials. Tf-idf was applied to the dataset, first with the entire feature list, followed by altered feature list with decreasing size to see if a more optimal set occurred within the entire list of features. The code used for the logistic regression trials can be found here: https://github.com/udel-biotm-lab/Chinese-Hamster-Ovary-Cell-Logistic-Regression.git.

#### Under-sampling

It is common practice to use under-sampling when the distribution of the sets is quite skewed, as in the cases of BP/non-BP (1049/8640 documents) and Glycosylation/non-Glycosylation BP (70/979 documents). We applied under-sampling of the majority class by varying the under-sampling rate with each iteration. The statistics of the overall efficacy of the model were taken with each trial. The fraction of the majority set was varied between 0.1 and 0.9 by intervals of 0.1. This means that the majority set, negative set, was cut down to 10% and all the way up to 90%. The test set and training set remained as specified above. As the under-sampling rate was adjusted, the feature set length was altered by 10% each time as well, thus, as the fraction of the negative set increased towards 90% by 10% intervals the feature set length increased at the same rate. Once under-sampling was conducted, logistic regression was run using the tf-idf.

## 3. Results and Discussion

### 3.1 LDA topic modeling

#### Comparative analysis of LDA topics and manual categories

We have compared the topics (“Topics”) uncovered by the LDA models with bioprocess categories (“Categories”) manually compiled in the CHO bibliome (6). The 15 human labeled categories and their document sizes are: Phenotype and Production Characteristics (“Phenotype”, 547 documents), Enzyme Analysis (“Enzyme”, 152), Glycosylation (“Glycosylation”, 70), Purification and Separation Methods (“Purification”, 55), Gene expression and Transcriptomics (“Transcriptomics”, 51), Modeling (“Modeling”, 36), Proteomics (“Proteomics”, 36), Metabolomics and Fluxomics (“Metabolomics”, 32), Metabolism and Metabolic Flux Analysis (“Metabolism”, 31), Expression and Transfection Methods (“Expression”, 30), Secretory Pathway and Product Secretion (“Secretion”, 29), Cell Line Construction and Characterization (“Cell Line”, 26), Genomics and Epigenetics (“Genomics”, 24), RNAs and codon usage (“RNAs”, 23 documents), and Culture Strategy and Bioreactor Design (“Culture”, 18). Note one remaining category in the CHO bioprocess bibliome, Review Articles or Other (116 documents) that consists of review articles on CHO cells and high-throughput data for CHO culturing, was presented as part of the “others” category in the results below.

The LDA model discovered 9 topics (Fig 2, S2 File) from the bioprocess documents. The top four topics, covering 20.5% to 12.6% of tokens (i.e., terms or words including bigrams and trigrams) of the corpus, account for a total of 65.5% of tokens in the whole corpus.

**Figure 2.**
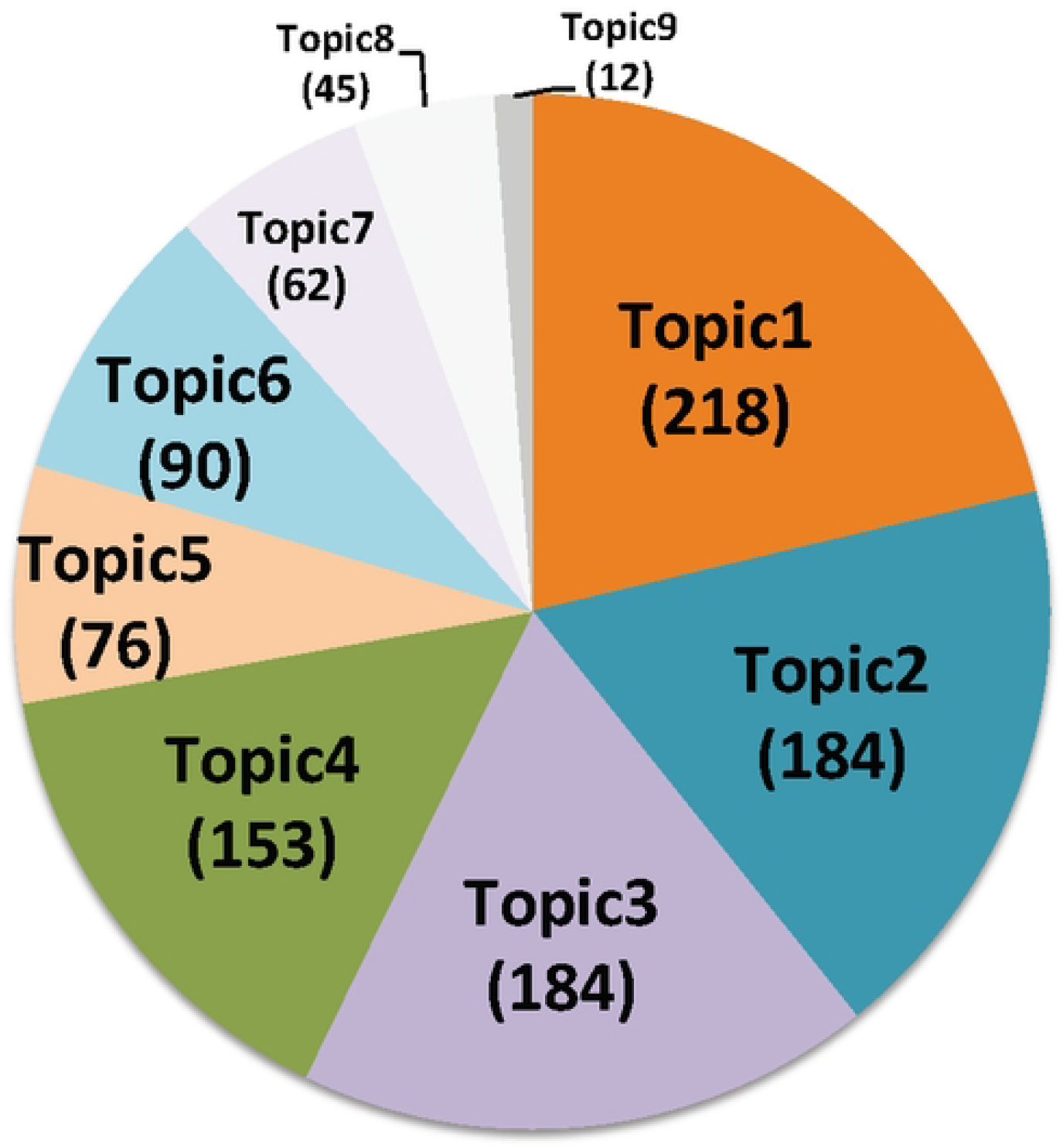
LDA topic categorization of CHO bibliome: distribution of bioprocessing documents in each of the 9 topics discovered by LDA model.

LDA allows multiple topics for each document, by showing the probability of each topic (10). For example, document for PMID 9043639 has probability of 0.49 for Topic-4 and 0.37 for Topic-1 according to the LDA model predictions (S2 Fig). To simplify further analysis, each BP document was assigned a representative Topic ID corresponding to the highest probability score (e.g., PMID 9043639 is assigned Topic-4 as its representative topic). To compare how LDA topics align with human category labels, heatmaps were generated where the columns show the human Category label and rows correspond to documents (PMIDs shown on the left) broken into different LDA topic groups (S3 Fig).

Fig 3 shows the comparative analysis of automatically generated LDA topics and the manually annotated categories. The overall distribution readily reveals that human category labels are differentially captured by LDA topics (Fig 3A). Among the four largest topics (containing 153 to 218 documents), Topic-1 is mapped to Categories “Phenotype”, “Transcriptomics”, “Proteomics”, and several other categories; Topic-2 to Categories “Phenotype”, “Expression”, “Cell Line”, “Secretion”; Topic-3 to “Phenotype”, “Glycosylation”, “Purification”, “Enzyme”; and Topic-4 to “Enzyme” and “Phenotype” (Fig 3B). While Topic-1, −2 and −3 spread over several topics, Topic-4 has only two major categories. The document sets for the remaining topics are much smaller and predominant with Category “Phenotype” (Fig 3A). Among the top 4 categories, “Phenotype”, “Enzyme”, “Glycosylation” and “Purification”, the largest is “Phenotype” which accounts for over 50% of all bioprocess publications in the CHO bibliome. Not surprising, it has a diverse distribution over many Topics (Fig 3C). In contrast, the other three categories all have only one dominant Topic each, “Enzyme” is dominant with Topic-4, and “Glycosylation” and “Purification” are dominant with Topic-3.

**Figure 3.**
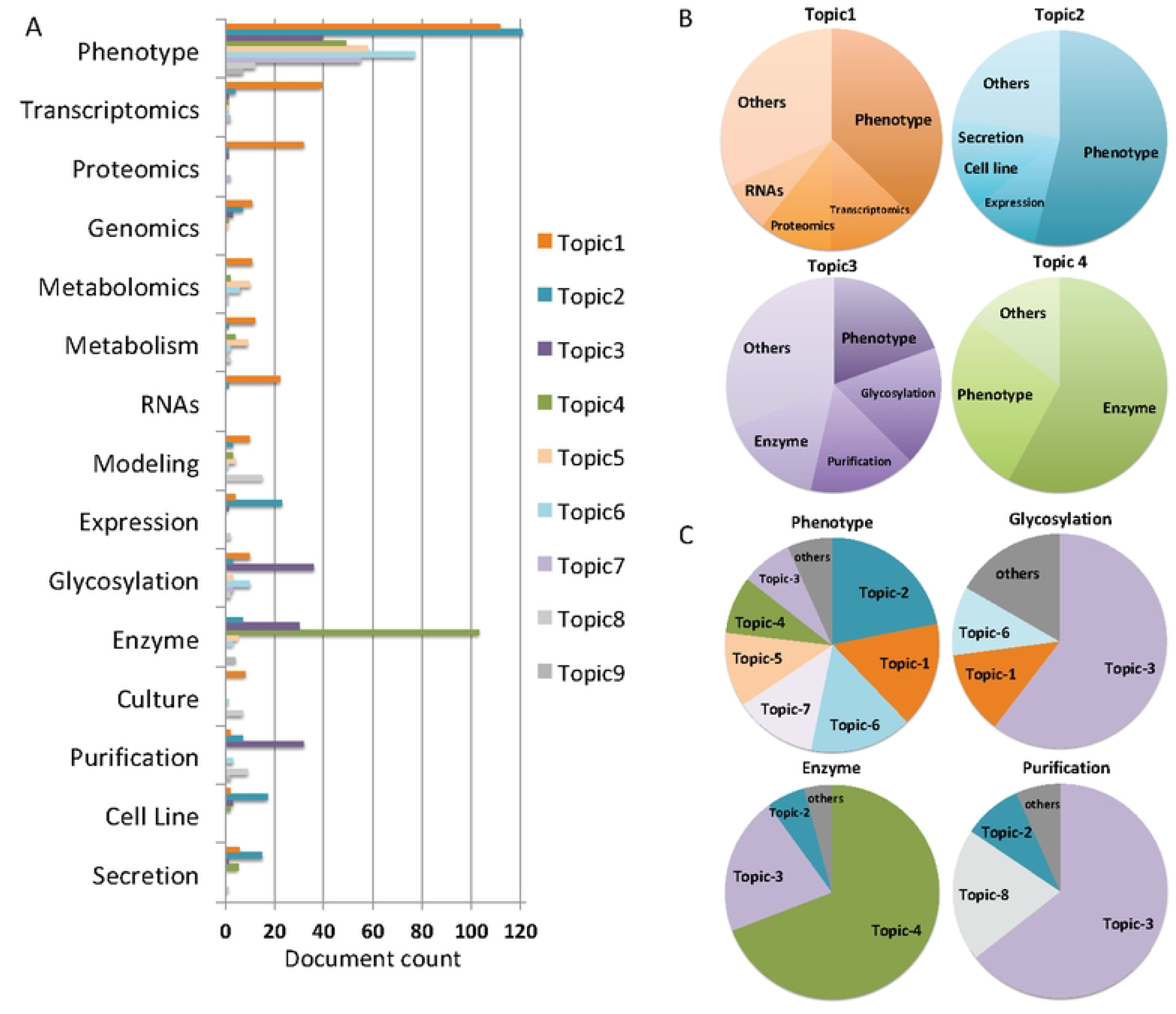
Comparison between automatically generated LDA topics and manually assigned categories. (A) Distribution of human-annotated categories among computer-generated LDA topics. (B) Distribution of the top four LDA topics in manual categories. (C) Distribution of the top four manual categories in LDA topics.

#### Interpretable terms in topic models

A basic question to ask about a topic model is whether the topics are interpretable to human. LDA represents documents as a mixture of topics, and a topic as a mixture of words, with different weights as the probability of those words appearing in the topic. Fig 4 shows the top terms for each topic and a pyLDAvis display from interactive topic model visualization (S2 File). In Fig 4A, each of the LDA topics is featured with top 30 most frequent terms with term weights (i.e., probabilities). Here, word “cell” is among the top 15 most frequent words for all topics. It is ranked first in Topic-4, −5, −6, −7, with the highest weight for Topic-4. On the contrary, word “mutant” is exclusive to Topic-4 among the top 30 words for all topics, therefore it is a discriminative key term in capturing a document into Topic-4. Fig 4B shows the pyLDAvis display of top 30 most frequent words for Topic-4. In addition to “mutant”, several terms such as “synthesis”, “biosynthesis”, “cholesterol”, “transport” have overlapping red bar and blue bar, indicating these terms are also frequent and exclusive to Topic-4 (also see PMID 9456320, 7742354, and 18946045 for examples, S3 File). With the terms mapped closely between Topic-4 and Category “Enzyme”, it is not surprising to see that majority of documents captured in Topic-4 are indeed in Category “Enzyme” in human annotation, and vice versa, indicating an intrinsic cohesiveness of human label and fitted LDA model for this topic (Fig 3A).

**Figure 4.**
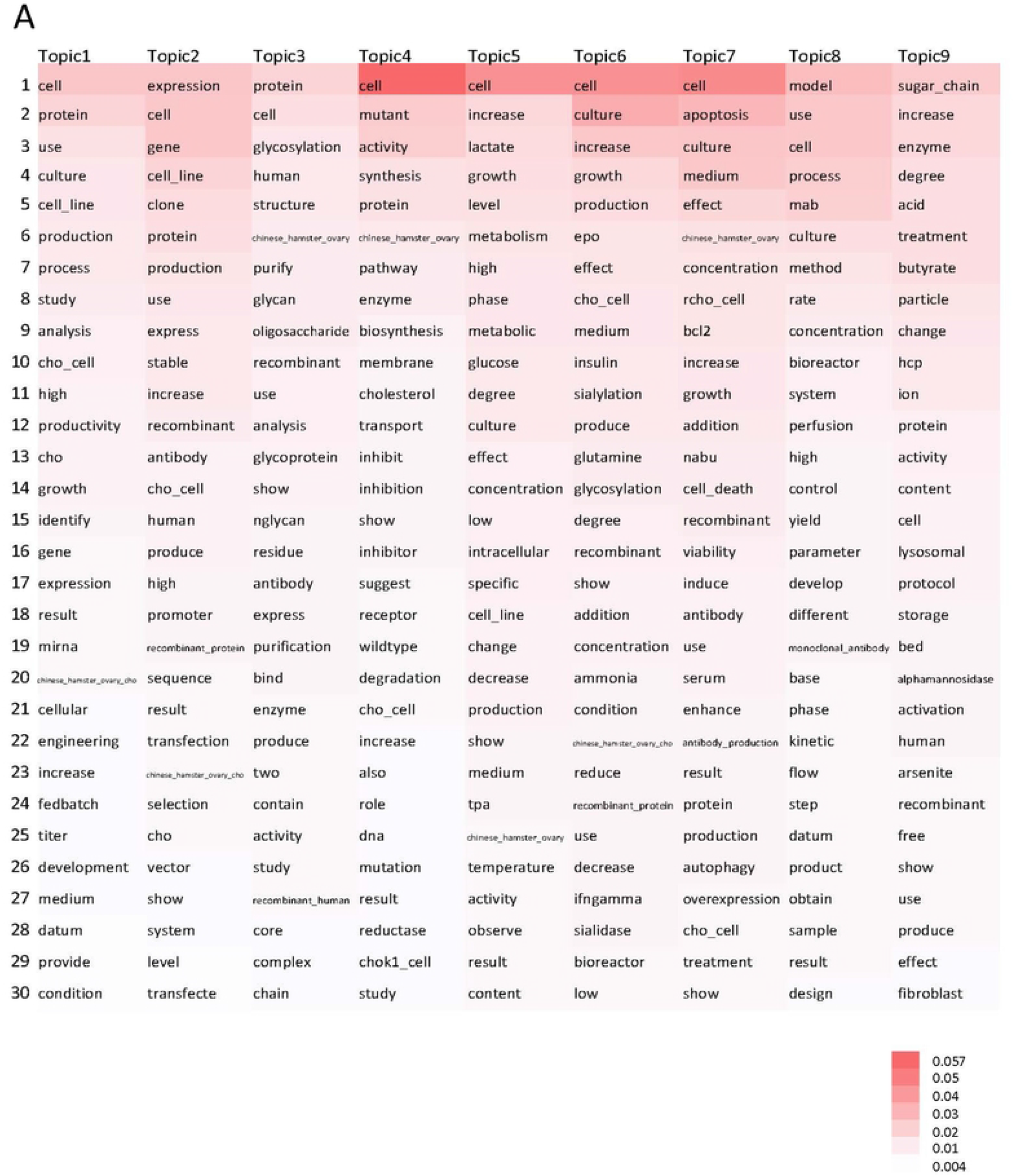
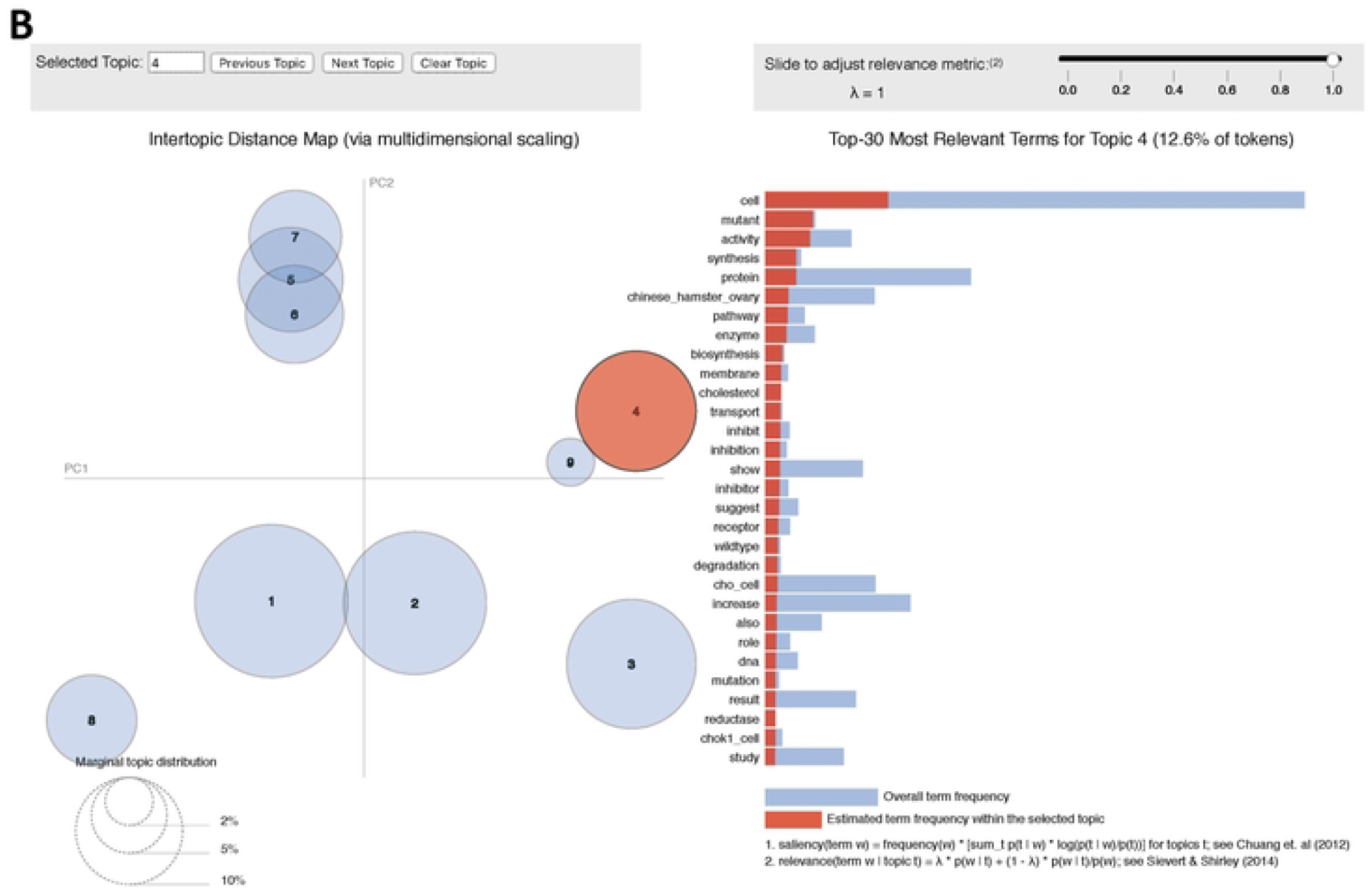
LDA topics with term probability. (A) The top 30 most frequent terms from nine LDA topics with weights. (B) Visualization of topic modeling results using pyLDAvis. Left shows semantic topic space, where each circle is a single topic and its size represents its importance in the model. The proximity between two circles reflects the semantic similarity of their concepts. Right shows Top-30 most salient terms for Topic-4. The terms (red bars) are in descending order of probability, and the blue bars show the terms’ frequency over whole corpus (i.e., a pair of overlaid bars represent both the corpus-wide frequency of a given term as well as the topic-specific frequency of the term). For a given term, when the red bar is almost the same length as the blue bar, it means it is a salient term almost exclusive to the topic.

In contrast, there are 3 significant human label categories for Topic-3, “Glycosylation”, “Purification” and “Enzyme” (excluding category “Phenotype” which have known intrinsically diverse documents). Among the most frequent words for Topic-3, “glycosylation”, “structure”, “purify”, “glycan”, “oligosaccharide”, “glycoprotein”, “nglycan”, “residue” are discriminative terms for categories “Glycosylation” and “Purification” (S3 File). Likewise the frequent and discriminative words for Topic-2 include “expression”, “gene”, “clone”, “stable”, “promoter”, “transfection”, “selection”, “vector”, which correlate well with categories “Expression”, “Cell Line” and “Secretion” where those words can be common and expected to occur together. In summary, our LDA model is able to cluster BP documents into topics with salient terms that are discriminative and descriptive for their underlying categories, and the computationally generated models correlate well with several human-labeled categories.

### 3.2 Logistic regression for classification

#### Binary representation

The results for binary representation of a term (presence or absence of a term in a document) serve as a baseline to see if each subsequent configuration improved the overall effectiveness of the classifier (Table 1). The Phenotype set obtained the best classification results. The BP set had lower F1-score, precision, and recall. The Glycosylation set had large fluctuations in the results and this could be the first sign that the small positive set and the large size difference between the positive and negative set is affecting the results of the classification.

**Table 1.** Logistic regression utilizing binary representation of terms.

| Document Set (Size) | F1-Score | Precision | Recall | Accuracy |
| --- | --- | --- | --- | --- |
| Phenotype/non-Phenotype (547/502) | 0.76 | 0.78 | 0.75 | 0.76 |
| BP/non-BP (1049/8640) | 0.61 | 0.66 | 0.57 | 0.92 |
| Glycosylation/non-Glycosylation (70/979) | 0.22 | 0.5 | 0.14 | 0.93 |

#### Term frequency-inverse document frequency (tf-idf)

Utilizing tf-idf, the length of the feature set can be seen. This allows feature engineering to find optimal configurations of the classifier. The df_min and df_max values were set to remove terms with very low occurrence rates. The results show that by using tf-idf the performance (especially precision) improved slightly for the BP vs. non-BP (Table 2). The statistics for logistic regression with a shortened feature list of top terms were quite similar to the trial run with all terms (Table 2). However, this configuration may be more desirable as the same results can be achieved with a much smaller set of terms.

**Table 2.** Logistic regression utilizing tf-idf with all terms and chosen top terms.

| Document Set | Feature Size | F1-Score | Precision | Recall | Accuracy |
| --- | --- | --- | --- | --- | --- |
| Phenotype/non-Phenotype | All terms (474) | 0.76 | 0.79 | 0.73 | 0.76 |
| BP/non-BP | All terms (706) | 0.62 | 0.75 | 0.53 | 0.93 |
| Phenotype/non-Phenotype | Top terms (250) | 0.75 | 0.76 | 0.75 | 0.75 |
| BP/on-BP | Top terms (500) | 0.63 | 0.75 | 0.54 | 0.93 |

For the Glycosylation set, the performance dropped to all zeros with a high accuracy level. This trend indicates that the model was predicting all of the documents as negatives and because the true number of positives was so limited the accuracy was 93%. This confirms the need for under-sampling. The accuracy is also much higher than the other statistics for the BP vs. non-BP which shows that under-sampling may be needed here as well.

#### Under-sampling

Previous results showed that the results were better for the phenotype/non- phenotype sets than with the other two classifications. We believe this is because the phenotype/non-phenotype document distribution is reasonably balanced, unlike the other two cases. We conducted under-sampling to address the unbalanced size of the positive and negative data for the Glycosylation/non-Glycosylation (70/972) and the BP/non-BP (1049/8640) sets. When the ratio of the positive to the negative data was set to be roughly 1:1, the performance was greatly improved (Table 3).

**Table 3.** Logistic regression utilizing under-sampling and tf-idf with chosen top terms (fraction of majority set = 0.1)

| Document Set | F1-Score | Precision | Recall | Accuracy |
| --- | --- | --- | --- | --- |
| Glycosylation/non-Glycosylation | 0.64 | 0.7 | 0.58 | 0.76 |
| BP/non-BP | 0.71 | 0.65 | 0.78 | 0.66 |

Table 2 shows that the best results were obtained in the case of phenotype/non-phenotype dataset. These results is based on the simple choice of representation of terms – whether or not the term appeared in the document. Table 3 shows the results of the same three classifications with use of tf-idf for the terms. We observed that this commonly-used representation of terms in information retrieval doesn’t have any significant impact on phenotype/non-phenotype classification but offers a slight improvement for BP/non-BP classification, with a significant gain in precision. This experiment also used a cut-off for the use of terms by applying a threshould for minimum and maximum document frequency (i.e., how many documents does a given term appear in). Further restrictions to terms by using top terms only did not show much change in the composite F1 scores.

We also noted that the BP/non-BP and Glycosylation/non-Glycosylation datasets are imbalanced with distribution being heavily skewed towards the negative set. This clearly impacted the results of these two classification tasks in contrast to the better balanced phenoype/non-phenotype. To address this situation, we applied under-sampling of the majority class and show in Table 3 the F1 scores improves significantly, especially in the gain of recall as is to be expected with undersampling the majority, negative class.

## 4. Conclusions

In this work, we have described approaches to analyze CHO cell literature that would be of general interest to researchers of CHO bioprocessing research and broader bioengineering and biotechnology community. It makes use of the existing CHO bibliome dataset previously manually labeled with research categories in 2016. Our unsupervised topic modeling enabled a detailed comparison between human labels and machine-generated topics, which empowers qualitative and quantitative understanding of the CHO literature set. Even though the size of the corpus for LDA is relative small in our current study from NLP perspective, select topics notably mirror some human manually assigned topic categories. With the insight gained from our LDA model, we further applied supervised learning for document classification to address the pressing need of classifying new unseen publications automatically instead of time-consuming manual labeling. Making use of terms as features for given topics, the effect of different feature representations on classifier performance is studied. Our study showcases important applications of text analytics on a biological scientific corpus: it discovers structural relations between topics and documents, summarizes corpus via visualization, and discusses challenges and future studies for consideration.

We have explored supervised deep learning method BioBERT (Bidirectional Encoder Representations from Transformers) due to its strength in classifying biomedical literature (20). We used the Google Cloud platform and conducted preliminary studies where the learning rate, epoch amount, and token limits, as well as other variables can be controlled (20, 21). This method is a possible path for supervised text classification of datasets such as the CHO bioprocess bibliome, but more research must be performed to test its applicability. BERT models are powerful models and tend to overfit when training data is not sufficiently large. This was a factor for not including their use in this work.

## 5. Acknowledgments

We thank CHO Genome to Phenome CHOg2p project community for their helpful discussions and suggestions, and appreicate Dr. Sarah Harcum (Clemson University) and Dr. Kelvin Lee (University of Delaware) groups for sharing their ideas at the early stage. This work is completed with the kind aid of Dr. Debarati Roy Chowdhury, Dr. Sachin Gavali, Dr. Cecilia Arighi, and Dr. Peng Su from University of Delaware.

## Supporting information

**S1 Figure. Overview of CHO bibliome bioprocessing set with manual categories**

**S2 Figure. Snip shot of Topic4Term.html in S3 File to show PMID 9043639**

**document with salient terms in color.**

**S3 Figure. Heatmap of documents by LDA topic-1 to −9 and manual categories.**

**S1 Table. Human labels of categories for CHO bioprocess set.**

**S1 File. Cross comparisons between LDA topics and manually assigned categories**

**for a subset with single dominant LDA topic assigned**.

**S2 File. LDA topics summarized with pyLDAvis in interactive html.**

**S3 File. Html documents by LDA topics with salient terms in color.**

